# NEMF inhibits proliferation and migration of ovarian cancer cells by blocking the PI3K/mTOR pathway

**DOI:** 10.1101/2024.11.20.624445

**Authors:** Qian Wang, Ying Zhu, Yilin Zhang, Chuangzhen Chen, Jing Fan, Aiqi Yin, Suiqun Guo, Yahui Liu

## Abstract

Nuclear export mediator factor (NEMF) is considered to be a tumor suppressor in Drosophila. Based on the analysis of data from an online database, NEMF is downregulated in ovarian cancer. However, the mechanism underlying its expression has not been reported thus far.In this study, the immunohistochemistry results showed that the expression of NEMF may be related to patient age and tissue differentiation. In vitro experiments indicated that overexpression of NEMF may inhibit cell proliferation and migration by inhibiting the PI3K/mTOR signaling pathway, thereby blocking cell cycle progression and promoting apoptosis (P < 0.05).Therefore, we speculate that NEMF is a potential tumor suppressor gene. Targeting NEMF may be an innovative therapeutic approach for ovarian cancer.

**Summary blurb:** NEMF is downregulated in ovarian cancer and the mechanism underlying its expression has not been reported thus far.In vitro experiments indicated that overexpression of NEMF inhibit cell proliferation and migration by inhibiting the PI3K/mTOR signaling pathway,thereby blocking cell cycle progression.

## Introduction

Ovarian cancer (OV) is associated with high mortality rates, posing a serious threat to women’s health and lives worldwide(L & Sr, 2020; M et al, 2022). According to data from the Global Cancer Research Agency published in 2022, the annual incidence rate of OV has reached 6.5/100,000 individuals, with approximately 320,000 new cases and 200,000 deaths recorded worldwide(F et al, 2024). The incidence of OV is second only to that of cervical cancer and endometrial cancer among female reproductive system tumors. However, it has the highest mortality rate among these tumors(A et al, 2022; Y et al, 2022). The early symptoms of this type of cancer are relatively insidious; thus, most patients are diagnosed at an advanced stage of the disease(Pa & Ua, 2023), resulting in a poor prognosis(Lheureux et al, 2019). Therefore, exploring the biological characteristics and potential therapeutic targets of OV is essential to improve the survival rates and quality of life of patients.

Studies have shown that cell signaling pathways play an important role in tumor development and occurrence(Sun et al, 2021), particularly the phosphatidylinositol 3 kinase/protein kinase B/mammalian target of rapamycin (PI3K/AKT/mTOR) signaling pathway(Jd et al, 2007). This pathway plays a crucial role in regulating cell proliferation, apoptosis, deformation, and migration(Ersahin et al, 2015; Glaviano et al, 2023). Moreover, its abnormal activation is closely related to the occurrence of numerous types of cancer, including OV(Stamp et al, 2016; Ediriweera et al, 2019; Nikolatou et al, 2023). PI3K regulates cell growth and proliferation by activating downstream AKT, which in turn affects mTOR(Tewari et al, 2022). The PI3K pathway is commonly upregulated in OV(Huang et al, 2020), and PI3K signaling is associated with improved survival and chemoresistance (Gasparri et al, 2017). Thus, therapeutic strategies targeting this pathway are becoming a hotspot in research on antitumor therapy.

Nuclear export mediator factor (NEMF) is a relatively newly recognized tumor-related gene. At present, there is limited knowledge regarding this gene. NEMF encodes a component of the ribosomal quality control complex(Bengtson & Joazeiro, 2010; Crowe-McAuliffe et al, 2021). The encoded protein contributes to the ubiquitin ligase listericin recognition and ubiquitination arrested 60S subunit(Thrun et al, 2021). Similar proteins in *Drosophila* act as tumor suppressors(X et al, 2005). The exact role of NEMF in cancer and the underlying mechanism remain unknown.

Therefore, the aim of this study was to preliminarily investigate the role of NEMF in the proliferation and migration of OV cells through the PI3K/mTOR pathway and elucidate the underlying molecular mechanism.

## Results

### OV DEG screening

Screening for OV DEGs was conducted using microarray datasets from TCGA database. The researchers initially standardized the mRNA profiles of OV samples and performed differential expression gene analysis using the limma software package in R language. A total of 18,521 DEGs were identified in this study (17,768 and 51 genes with high and low confidence, respectively) (Fig 1A). These DEGs are key indicators of molecular alterations in OV, potentially highlighting key pathways and mechanisms involved in disease occurrence and progression. In addition, bioinformatic methods can be used to identify key DEGs and explore their biological expression patterns in OV. By constructing PPI networks and conducting functional enrichment analysis, the specific roles of these genes in cancer occurrence can be revealed, providing new therapeutic strategies and diagnostic markers for the effective treatment of OV.

**Figure 1.**
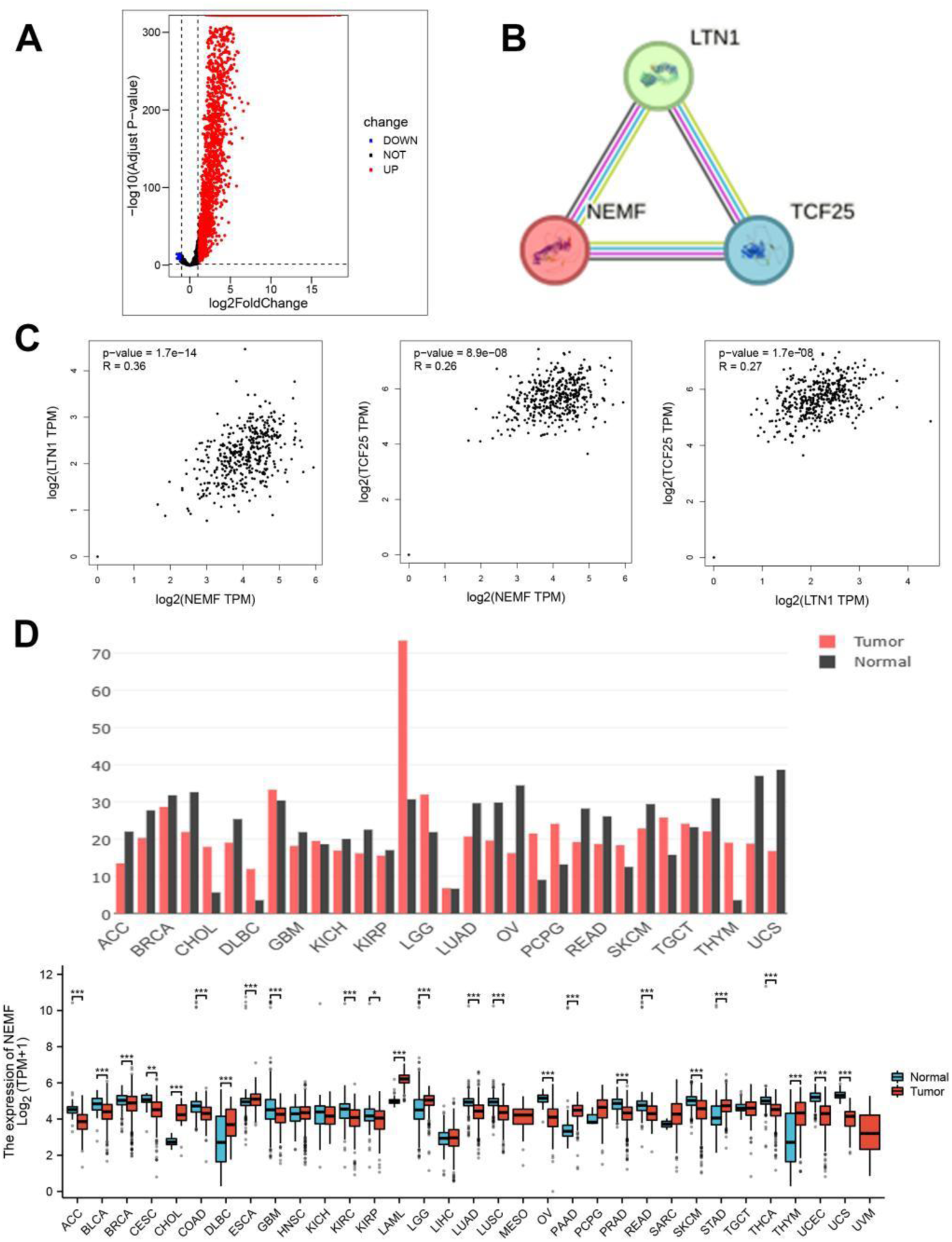
Volcano plots, interaction, and pan-cancer analysis of genes. **(A)** Volcano plots of DEGs observed in the difference analysis dataset. **(B)** PPI network was constructed, leading to the identification of three pivotal hub genes. **(C)** Expression levels of NEMF, LTN1, and TCF25. **(D)** Expression of NEMF mRNA at the pan-cancer level. NEMF mRNA expression levels in 33 human tumor tissues and corresponding normal tissues from the GEPIA database. \**P* < 0.05, \*\**P* < 0.01, \*\*\**P* < 0.001, \*\*\*\**P* < 0.0001.

We analyzed a recognition module of central gene selection based on cancer cell interactions. Three genes with downregulated expression were identified as central genes by STRING enrichment, namely NEMF, listerine E3 ubiquitin protein ligase 1 (LTN1), and transcription factor 25 (TCF25) (Fig 1B). We found that NEMF, LTNI, and TCF25 genes are lowly expressed and have an interactive relationship in OV. In addition, these genes play key roles in protein synthesis and quality control.

The identification and investigation of these genes may assist in understanding the biological mechanisms of OV and developing new therapeutics.

Furthermore, we used linear regression to observe the expression levels of NEMF and the correlation with LTN1 and TCF25. We found that the coefficient of genetic correlation of NEMF with LTN1 and TCF25 in the ovary was 0.36 and 0.26, respectively. The coefficient for the correlation between TCF25 and LTN1 was 0.27 (Fig 1C).

### Pan-cancer analysis and gene interaction

In the GEPIA database, we found that NEMF was expressed in all tissues. Analysis of TCGA tumors and adjacent normal tissues revealed that NEMF expression was significantly downregulated in 18 cancer types (i.e., adrenal cortical carcinoma, bladder urothelial carcinoma, breast invasive carcinoma, cervical squamous cell carcinoma and endometrial adenocarcinoma, colorectal adenocarcinoma, glioblastoma multiforme, chromophoric cell carcinoma of the kidney, clear cell carcinoma of the kidney, and renal papillary cell carcinoma, lung adenocarcinoma, lung squamous cell carcinoma, ovarian serous cystadenocarcinoma, prostate adenocarcinoma, rectal adenocarcinoma, skin melanoma, thyroid cancer, uterine endometrial carcinoma, and uterine carcinosarcoma). In contrast, NEMF expression was upregulated in 12 types of cancer (i.e., cholangiocarcinoma, lymphoid tumor diffuse large B-cell lymphoma, esophageal cancer, head-and-neck squamous cell carcinoma, acute myeloid leukemia, low-grade glioma of the brain, pancreatic adenocarcinoma, pheochromocytoma and paraganglioma, sarcoma, gastric adenocarcinoma, testicular germ-cell tumor, and thymoma) (Fig 1D).

### NEMF mRNA expression and infiltration analysis in OV

To examine NEMF expression in OV progression, the mRNA and protein levels in OV samples and normal tissue samples were studied using data from publicly available datasets and clinical samples. We compared gene expression levels in OV in the GEPIA database, and found that NEMF expression was downregulated (Fig 2A). GO analysis indicated that the main biological processes associated with feature gene enrichment were functions related to neuron cell body, synaptic organization, small GTPase signal transduction, and protein localization (Fig 2B).

**Figure 2.**
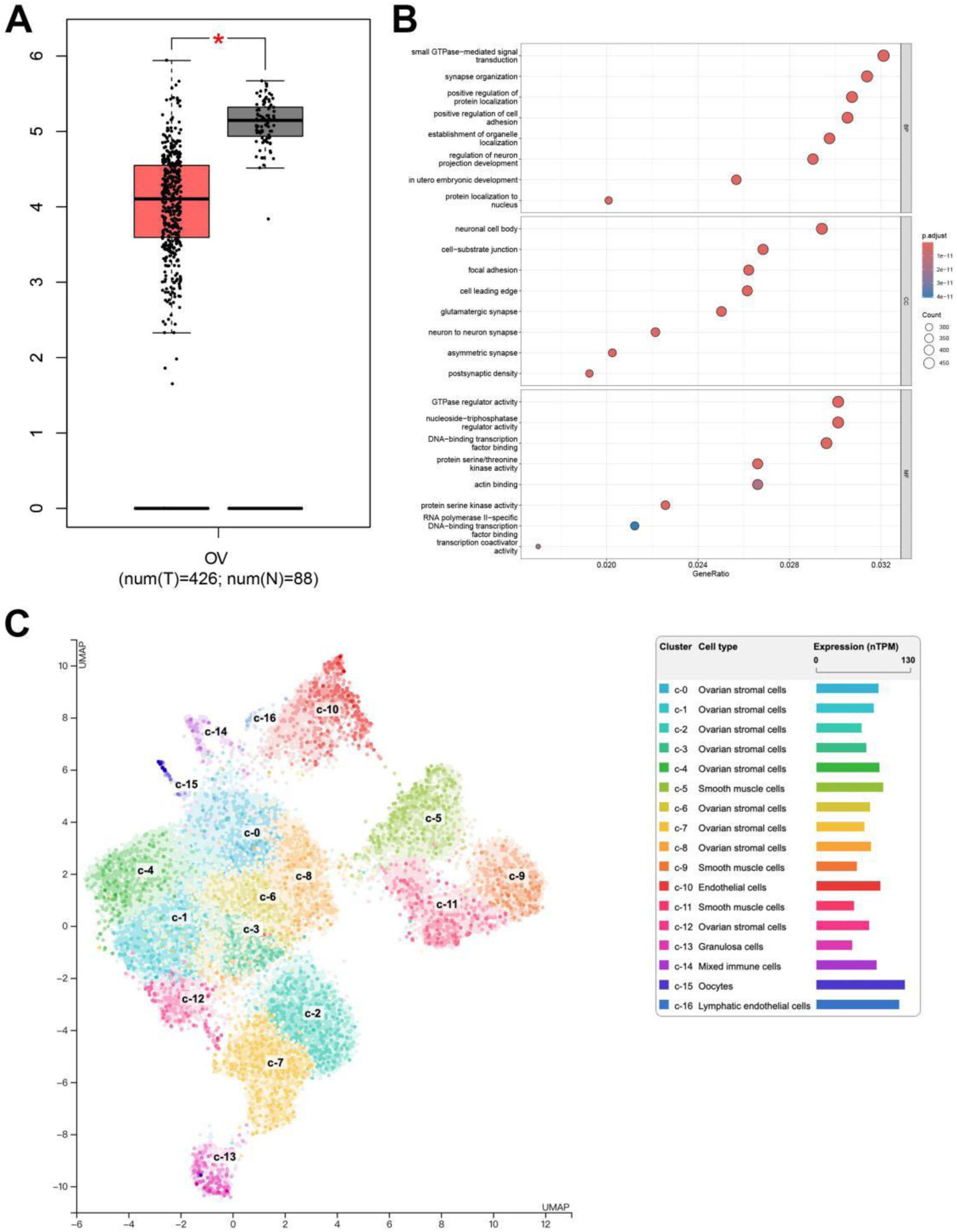
Enrichment analysis of the NEMF gene. **(A)** Differences in the expression of NEMF in OV tissues and corresponding normal tissues from the GEPIA database. **(B)** NEMF-related CC, BP, and MF items. **(C)** Distribution of different cell types in the sample.

In the distribution of different cell types in the sample, we found that c-0, c-1, c-2, c-3, c-6, c-7, and c-8 were ovarian stromal cells, while c-9, c-10, and c-11 were smooth muscle cells. Overall, the expression levels of ovarian stromal cells and some smooth muscle cells were high, whereas those of oocytes, lymphoendothelial cells and granulosa cells were low (Fig 2C).

### Relationship between NEMF expression and tumor-infiltrating immune cells

Since tumor-infiltrating immune cells are associated with OV prognosis, we performed an analysis to determine whether NEMF expression is associated with immune invasion in OV. Analysis of immune infiltration was performed according to NEMF expression (Fig 3A). To ensure the accuracy of the results, we also used TIMER to examine the relationship between NEMF expression and immune infiltration levels in different grades of OV. Assessments were performed in all OV samples to explore the association between key DEGs and infiltrating immune cells. In OV samples, NEMF, LTN1, and TCF25 exhibited clear patterns of association between genes and various types of infiltrating immune cells. A negative association was found between NEMF and B cells, CD8+ T cells, CD4+ T cells, macrophages, neutrophils, and dendritic cells. LTN1 was negatively correlated with B cells, CD8+ T cells, neutrophils, and dendritic cells; in contrast, it was positively correlated with CD4+ T cells and macrophages. B cells, CD8+ T cells, neutrophils, and dendritic cells demonstrated positive correlations with TCF25, while CD4+ T cells and macrophages displayed negative correlations. The data revealed the presence of a complex interrelationship in this gene–immune cell network (Fig 3B).

**Figure 3.**
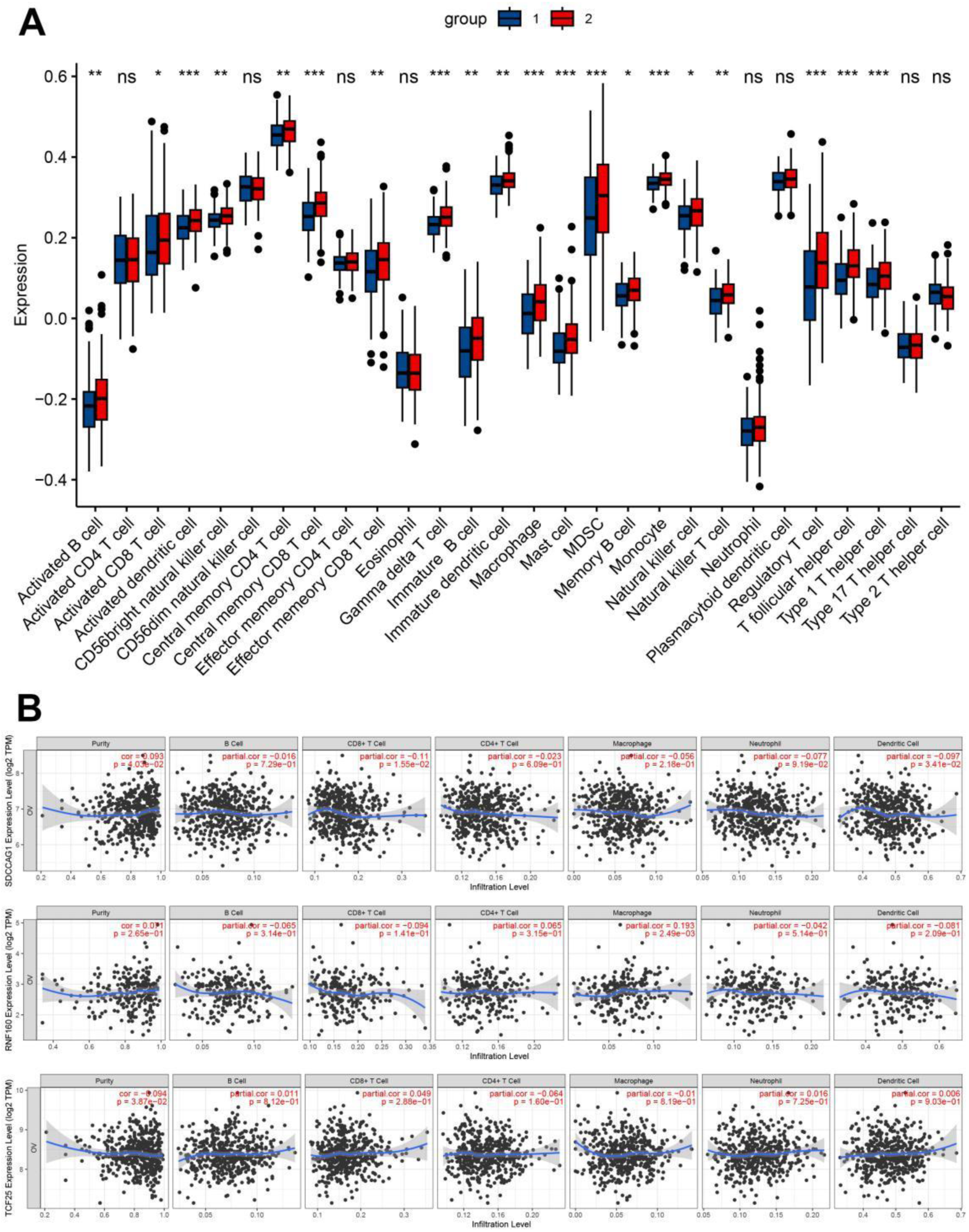
Tumor invasion was correlated with NEMF expression. **(A)** Expression of NEMF according to the level of immune infiltration. **(B)** NEMF expression in OV was positively correlated with infiltration of CD8 T cells, T cells, macrophages, column cells, MDSC, and monocytes.

### OV subcellular localization and decreased NEMF levels were associated with adverse outcomes

The subcellular localization of NEMF in OV cells was studied using the HPA database. The NEMF gene encodes A protein that is present in the cytoplasm and nucleosome (Fig 4A). The results of the overall survival and progression-free survival analysis showed that upregulation of NEMF was positively associated with the survival of patients with OV. Patients with high NEMF expression had longer overall survival (*P* = 0.036, hazard ratio = 0.79) (Fig 4B) and progression-free survival (*P* = 0.036, hazard ratio = 0.80) (Fig 4C) than those with lower NEMF expression. These results indicate that NEMF may be useful for predicting the survival of patients with OV.

**Figure 4.**
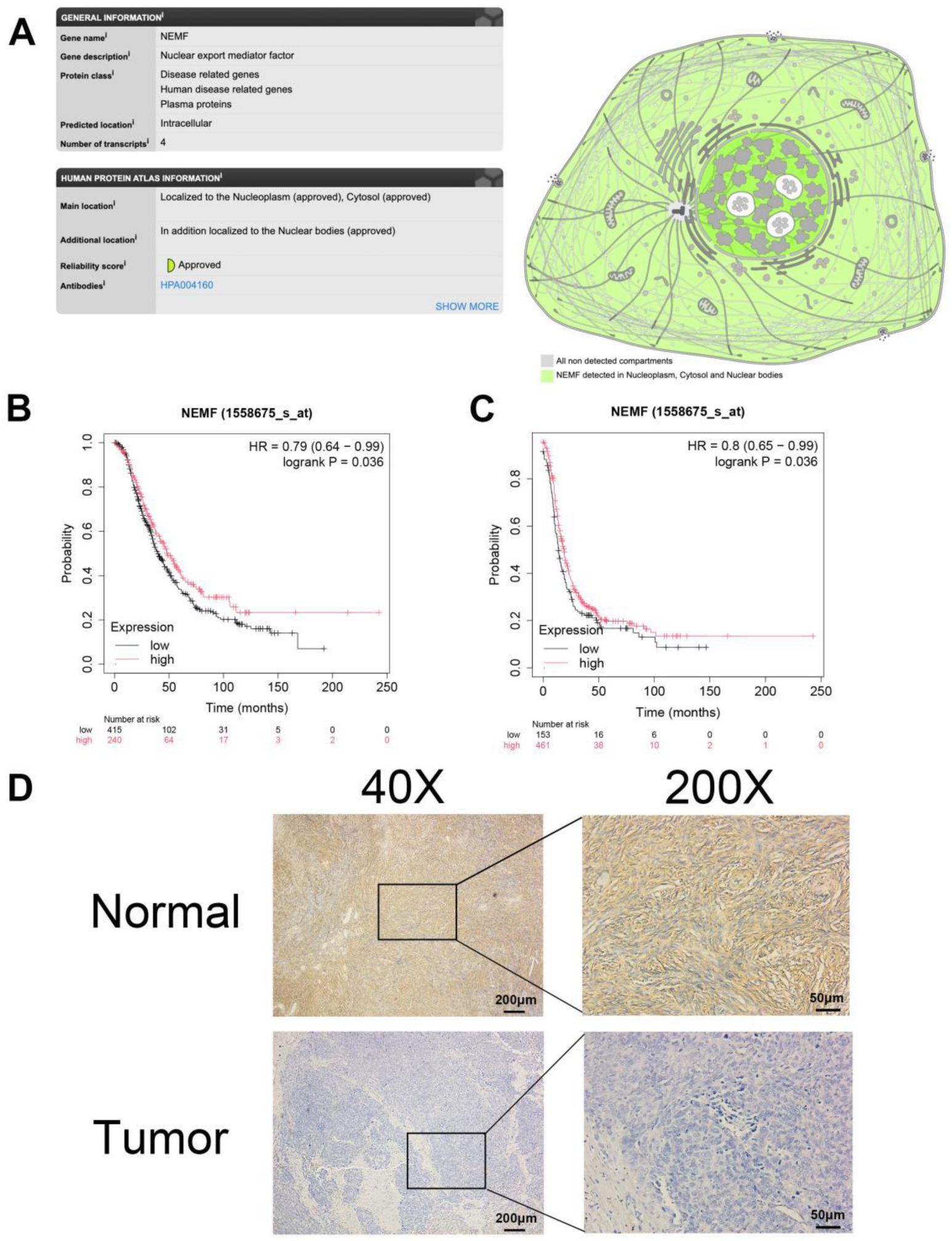
Expression and prognostic value of NEMF protein in OV. **(A)** Subcellular localization of NEMF protein. **(B)** OS curve of high/low NEMF expression group in OV. **(C)** PFS curve of high/low NEMF expression group in OV. **(D)** Immunohistochemical staining. The expression of NEMF was higher in normal tissues versus OV tissues.

Immunohistochemical staining was performed to further confirm the protein levels of NEMF. NEMF protein expression was significantly downregulated in OV tissues compared with normal tissues (Fig 4D). Chi-squared test and univariate Cox proportional hazards analysis showed that the expression of NEMF may be related to age and tissue differentiation (Tables 1 and 2). In summary, the present findings demonstrated that NEMF expression is reduced in OV and may be associated with poor prognosis.

**Table 1.**
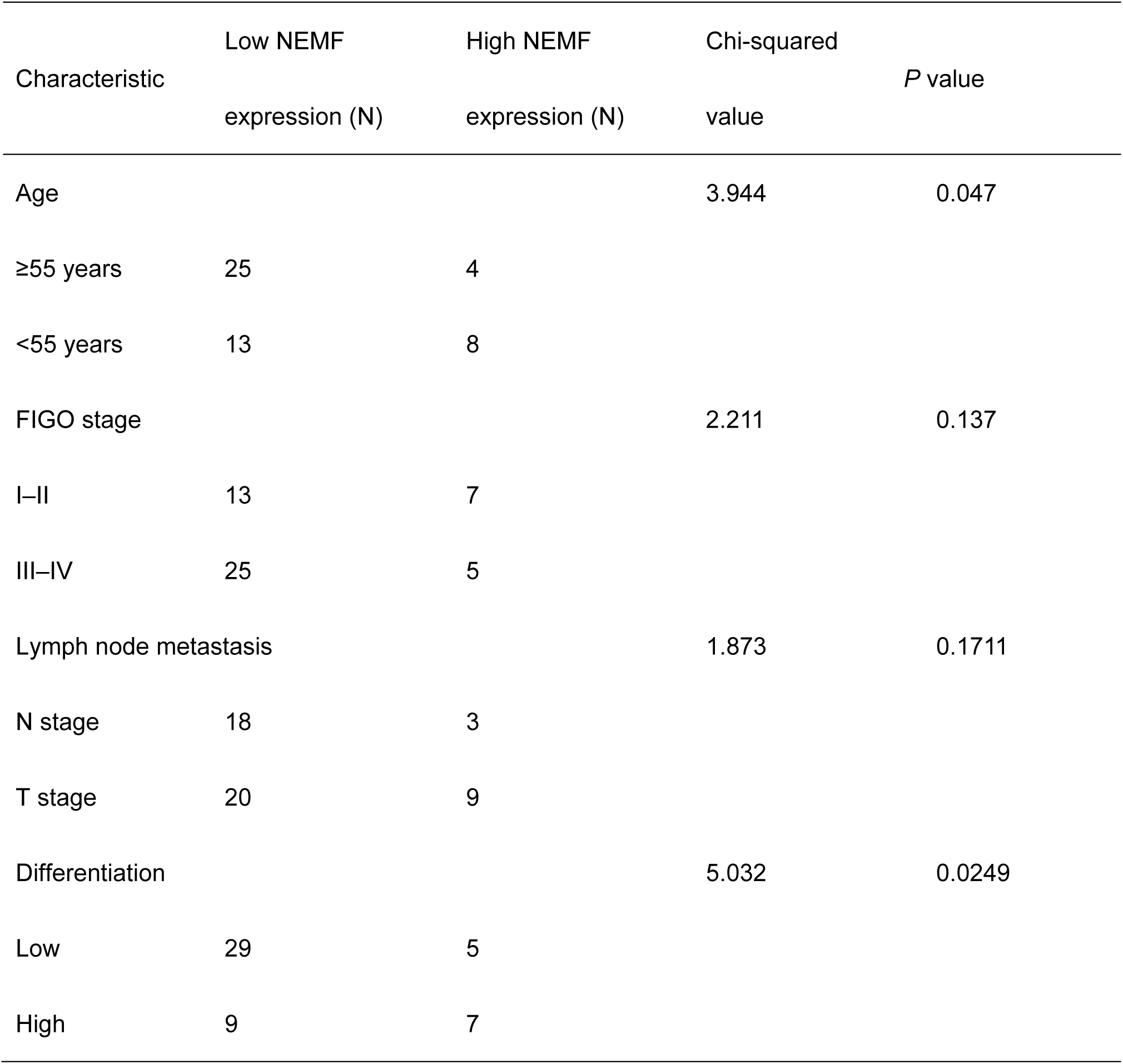
Correlation of NEMF expression in OV with clinicopathologic characteristics.

**Table 2.**
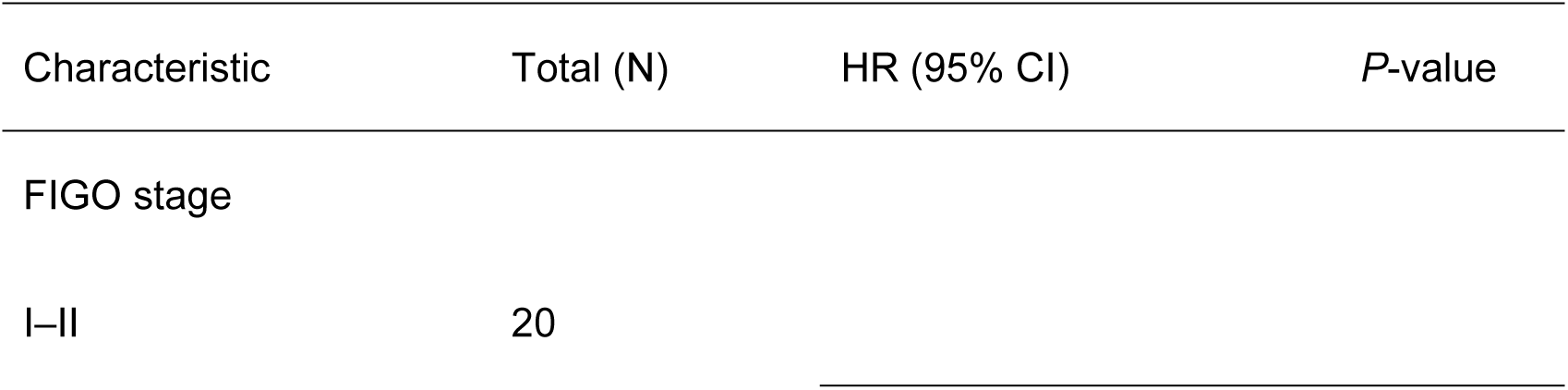
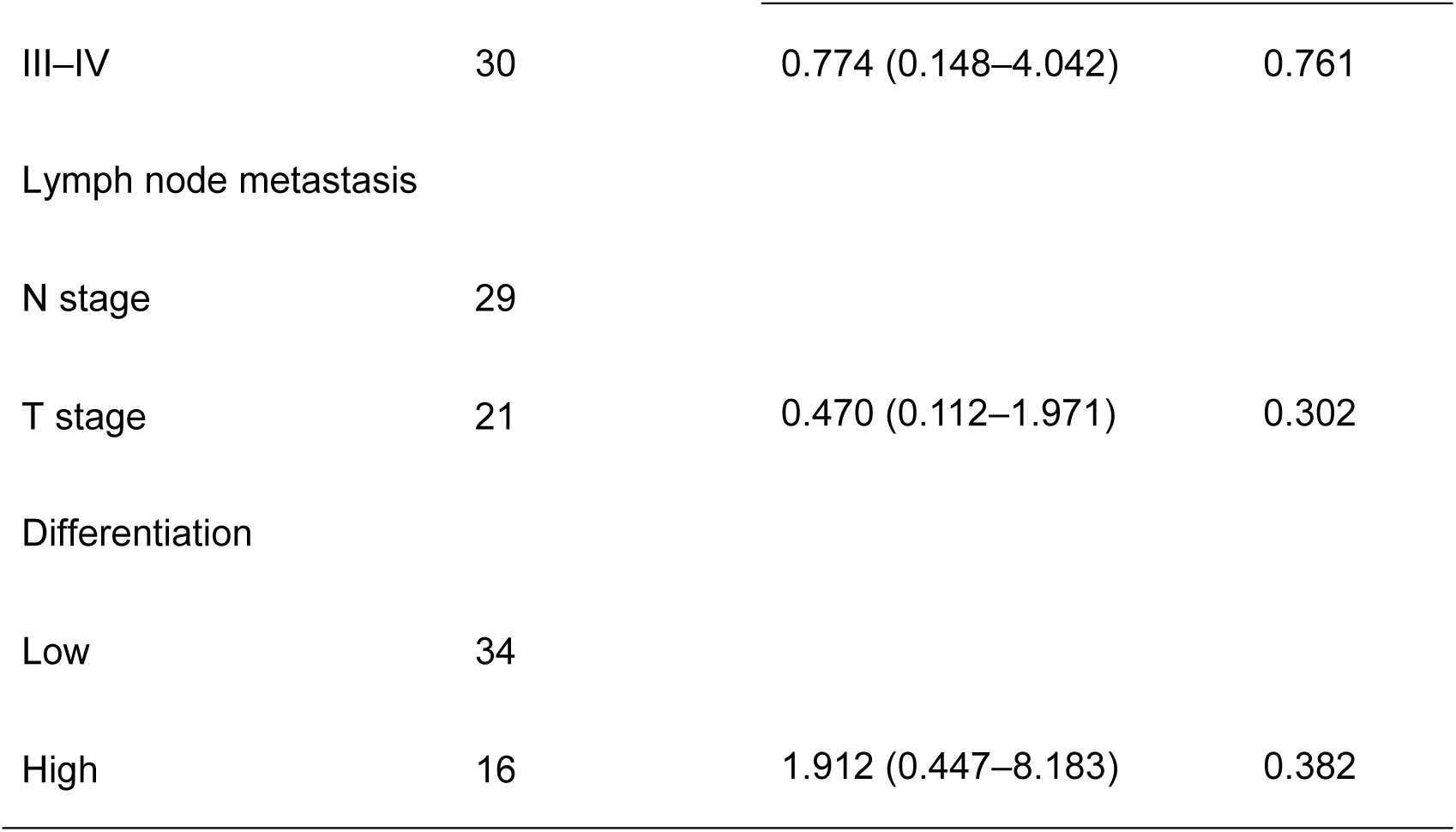
Analysis of risk factors affecting the prognosis of OV based on the univariate Cox proportional hazards model.

### OE of NEMF inhibited the proliferation and migration of OV cells

We first verified the basal expression of NEMF at the protein level. NEMF expression was significantly higher in normal ovarian epithelial cells (IOSE80) than in OV cell lines (OVCAR3 and SKOV3). This suggests that NEMF plays an important role in the physiological function of the ovary, and its decreased expression in OV cells may be related to the occurrence and development of tumors. These results were consistent with those obtained from the bioinformatics analysis (Fig 5A).

**Figure 5.**
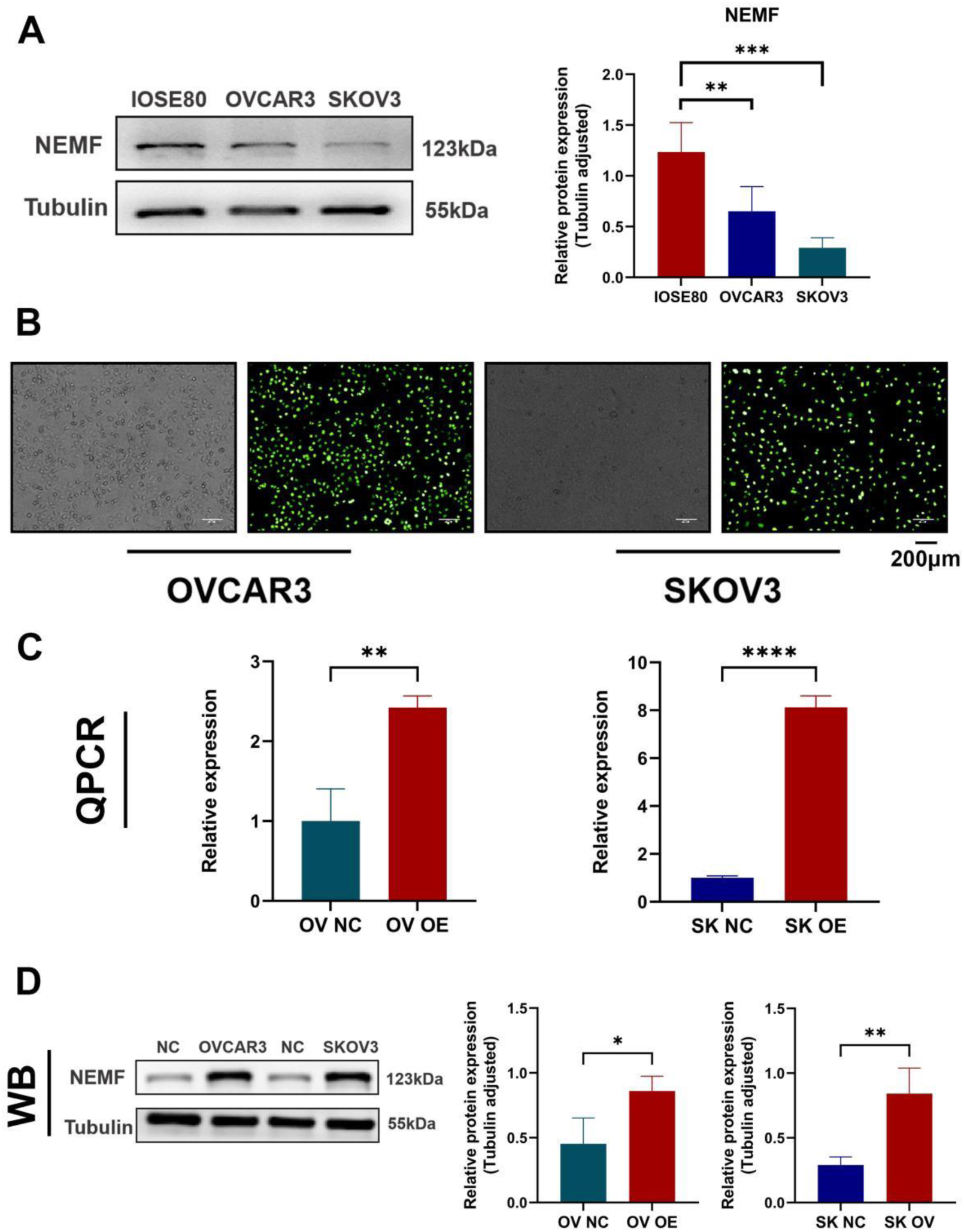
Expression levels of NEMF. **(A)** Basal expression of NEMF at the protein level. **(B)** Green fluorescence transfection efficiency. **(C)** NEMF overexpression validated by WB and quantitative PCR. \**P* < 0.05, \*\**P* < 0.01,\*\*\**P* < 0.001,\*\*\*\**P* < 0.0001.

To investigate the mechanism through which NEMF affects the proliferation and migration of OV cells, we constructed NEMF-overexpressing viruses and cultured OVCAR3 and SKOV3 cell lines with stable OE of NEMF. The transfection efficiency of GFP was observed and quantified by fluorescence microscopy (Fig 5B). Next, we investigated the phenotype by formal validation of OE. The efficiency was verified by quantitative PCR and western blotting (Fig 5C). NEMF expression was significantly higher in the OE group versus the negative control (NC) group, indicating that the OV cell lines (OVCAR3 and SKOV3) were successfully and stably transfected with the NEMF-overexpressing virus.

After verifying the OE efficiency, we investigated the proliferation and migration of OV cells with OE of NEMF *in vitro*. Firstly, CCK-8 assay results showed that the optical density values at 24, 48, 72, and 96 h were higher in the NC group versus the OE group in both OVCAR3 and SKOV3 cells (Fig 6A). Secondly, the EdU staining results showed that the proportion of OVCAR3 and SKOV3 cells in the proliferative phase was lower in the OE group than in the NC group (Fig 6B). The plate cloning experiment also yielded consistent results. In OVCAR3 and SKOV3 cells, fewer and smaller cell clones were formed in the OE group versus the NC group (Fig 6C). The above results indicate that OE of NEMF can inhibit OV cell proliferation.

**Figure 6.**
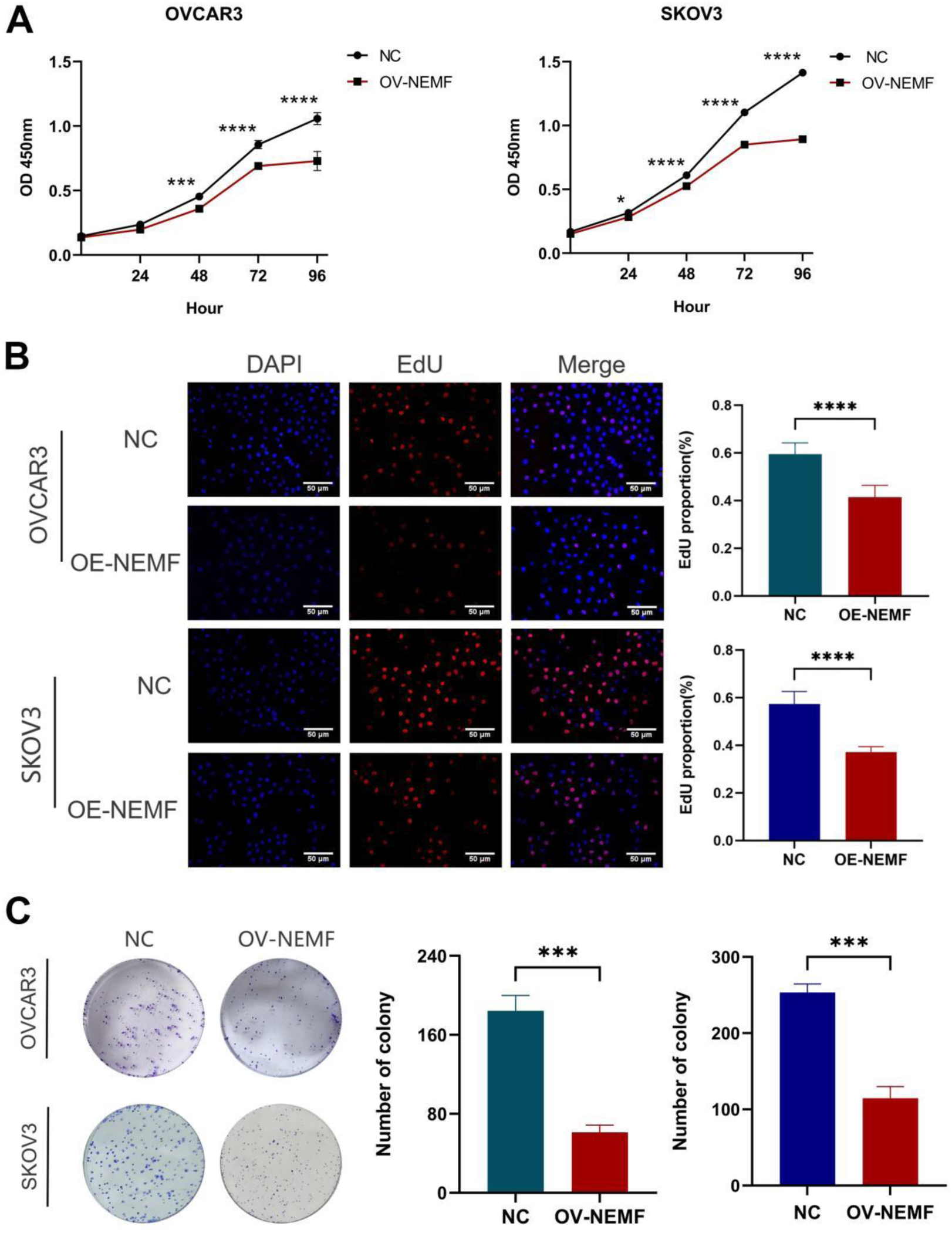
NEMF inhibited the proliferation of OVCAR3 and SKOV3 cells *in vivo*. **(A)** CCK-8 assay was used to detect the OD value of OV cells for 5 consecutive days. **(B)** EdU proliferation assay was used to detect cells in the proliferation phase in the NC and OE groups. **(C)** The number and size of cell clones in the NC and OE groups were measured by plate cloning after 7 days. \*\*\**P* < 0.001, \*\*\*\**P* < 0.0001.

In addition, wound healing experiments showed that the healing ability of cells was significantly weakened in the OE group compared with the NC group (Fig 7A). The ghrocyclic peptide staining experiments also showed changes in the cell morphology and a reduction in the number of microfilaments on the cytoskeleton in the OE group compared with the NC group (Fig 7B). Results of western blotting analysis of cell migration proteins indicated that the NC group expressed higher levels of N-cadherin protein and lower levels of E-cadherin protein compared with the OE group (Fig 7C). The above results showed that OE of NEMF inhibited the migratory ability of cells. We sought to further understand the specific mechanisms through which NEMF affects OV cell proliferation. Thus, we used flow cytometry to study the cell cycle of OV cells after OE of NEMF. Compared with the NC group, the OE group exhibited a higher proportion of cells in the G1 phase and a lower proportion of cells in the G2 and S phases following OE of NEMF (Fig 8A). The results showed that OE of NEMF prevented the cell cycle transition from the G1 phase to the S phase, resulting in cell arrest in the G1 phase.

**Figure 7.**
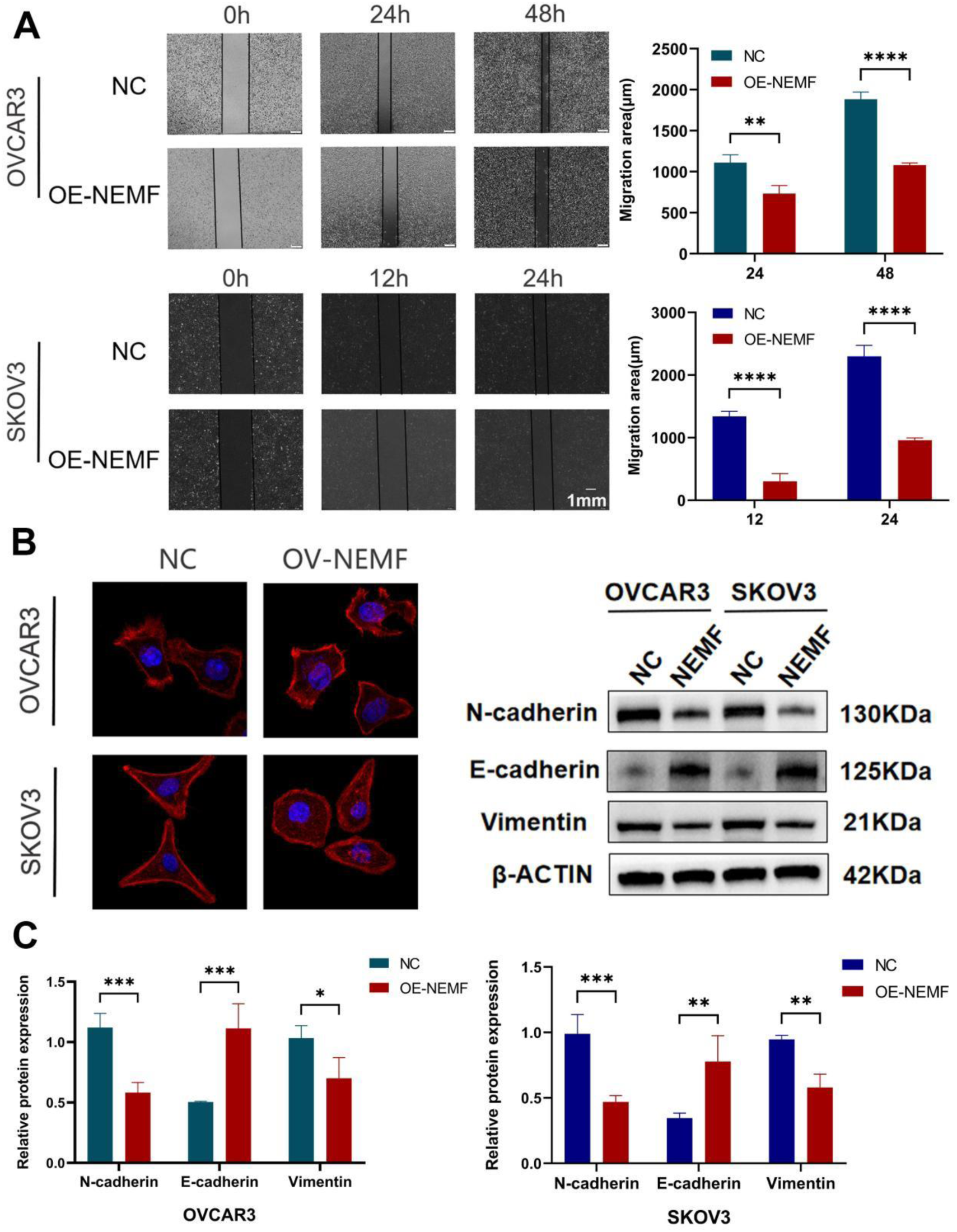
NEMF inhibited the migration of OVCAR3 and SKOV3 cells *in vivo*. **(A)** A wound healing experiment was conducted to compare the distance of migration between the NC and OE groups.**(B)** The migratory ability of cells was detected by ghrocyclic peptide staining experiments. **(C)** Expression results for proteins related to cell migration. \**P* < 0.05, \*\**P* < 0.01,\*\*\**P* < 0.001, \*\*\*\**P* < 0.0001.

**Figure 8.**
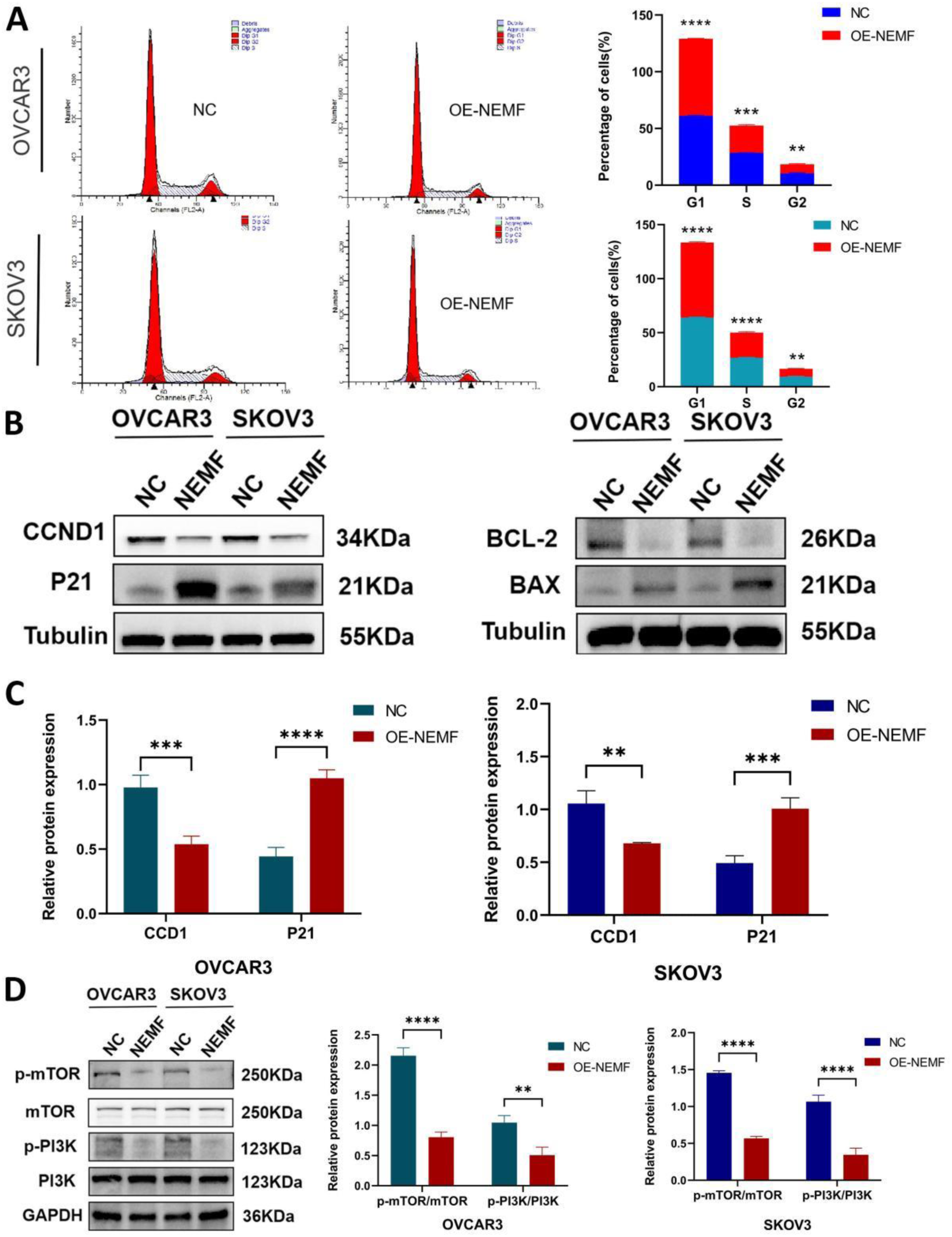
Results of the cell cycle, apoptosis, and PI3K/mTOR pathway analyses. (A) The cell cycle in the NC and OE groups was detected by flow cytometry. **(B)** Cell cycle and expression of apoptosis-related proteins. **(C)** Expression of proteins related to the PI3K/mTOR signaling pathway \*\**P* < 0.01,\*\*\**P* < 0.001, \*\*\*\**P* < 0.0001.

Western blotting of cell cycle-associated proteins revealed lower levels of CCND1 in the NC group versus the OE group, and higher levels of P21 protein in the NC group versus the OVCAR3 and SKOV3 cell lines (Fig 8B). It is well established that CCND1 promotes the transition from the G1 to the S phase, whereas P21 inhibits this transition. The above results revealed that OE of NEMF may arrest the cell cycle of OV cells in the G21 phase by inhibiting CCND1 and promoting the expression of P21. Western blotting results of apoptosis-related proteins showed that the expression of BCL-2 protein in the NC group was higher than that recorded in the OE group.

Moreover, the expression of BAX protein was lower in the NC group versus the OE group (Fig 8B). This evidence indicates that NEMF may promote apoptosis in OV cells.

Finally, according to previous data, it is well established that the PI3K/mTOR signaling pathway is often activated in OV and plays an important role in cell cycle and apoptosis. Therefore, we investigated whether OE of NEMF exerts an effect on the PI3K/mTOR signaling pathway. Results from western blotting revealed significantly lower levels of PI3K/mTOR signaling pathway signature proteins (i.e., p-PI3K/PI3K and p-mTOR/mTOR) in the OE group compared with the NC group (Fig 8C), suggesting that OE of NEMF may inhibit the PI3K/mTOR signaling pathway.

Based on the above findings, OE of NEMF may affect the expression of proteins involved in the cell cycle and apoptosis by inhibiting PI3K/mTOR signaling. These effects lead to cell cycle arrest in the G1 phase, promote apoptosis, and eventually inhibit the malignant biological behavior of OV cells.

## Discussion

OV is a common type of malignant gynecological tumors. The vast majority of ovarian tumors originate from the epithelium (90%)(Dh & Wx, 2013), including four subtypes (i.e., serous adenocarcinoma, clear cell carcinoma, endometrial adenocarcinoma, and mucinous adenocarcinoma)(Liu & Zheng, 2011; Duska & Kohn, 2017). Because the early clinical symptoms of OV are not obvious and non-specific, most patients are diagnosed at a clinically advanced stage of the disease, thereby missing the opportunity for radical cure(Akter et al, 2022; Chen et al, 2022; X et al, 2024). Studies have shown that approximately 75% of patients with OV are clinically diagnosed at an advanced stage due to the lack of early symptoms(Siegel et al, 2021; Chen et al, 2023). At present, the first-line treatment of OV involves cytoreductive surgery combined with platinum-based chemotherapy(Vaughan et al, 2011; Jiang et al, 2023). However, most patients with OV eventually develop platinum resistance and experience tumor recurrence(Godbole et al, 2023; Del Bufalo & Damia, 2024; Miras et al, 2024; Nunes et al, 2024). Consequently, the overall 5-year survival rate of these patients is only 30–40%(Solidoro et al, 2024).

NEMF is an essential component of the ribosome quality control complex (RQC). It extracts partially synthesized nascent chains from stalled ribosomes and degrades them through ubiquitin-mediated proteasome degradation (Shao et al, 2015; Hickey et al, 2020; T et al, 2021; Thrun et al, 2021). It has been reported that NEMF may be associated with neuromuscular diseases(Pb et al, 2020), and its mutations cause damage to the central nervous system(A et al, 2021). The present study is the first to investigate the mechanism underlying the effects of NEMF in cancer.

Initially, we used online databases such as TCGA, GEPIA, Kaplan–Meier plotter and R language to carry out basic expression analysis, pan-cancer analysis, enrichment analysis, and prediction of survival and prognosis in relation to NEMF. The results showed that NEMF was downregulated in OV, and its low expression was closely related to the poor prognosis of OV. Based on OV cell lines and clinical samples, we performed *in vitro* experiments to verify the reliability of the database. The experimental results showed that the protein levels of NEMF in OV cell lines were significantly reduced compared with those recorded in normal cells. Immunohistochemical experiments also confirmed the low expression of NEMF in OV and its prognostic significance at the tissue level.

In addition, we conducted a series of cell function experiments to investigate the effect of NEMF on the malignant biological behavior of OV cells. These experimental results showed that NEMF plays an inhibitory role in the proliferation and migration of OV cells. According to a number of studies, tumor cells rely heavily on the PI3K/mTOR signaling pathway for growth, proliferation, and survival(Huang et al, 2022; L et al, 2022), and this pathway is closely related to the cell cycle(Cao et al, 2023; Occhiuzzi et al, 2023). In the NEMF OE group, we observed changes in the cell cycle and the expression of apoptosis-related proteins (i.e., CCND1, P21, BCL-2, BAX)(K, 2022; Knudsen et al, 2022; Kaloni et al, 2023; N et al, 2023), indicating that NEMF can inhibit the cell cycle and apoptosis. However, the phosphorylation levels of key molecules of the PI3K/mTOR pathway, such as p-PI3K and p-mTOR, decreased significantly, denoting that NEMF can inhibit the activity of this signaling pathway.

Taken together, the results of the present study showed a decrease in the expression of NEMF in OV. Low expression of NEMF is associated with poor prognosis. It was demonstrated that NEMF suppresses the malignant behavior of OV cells. Additionally, OE of NEMF in OV cells may arrest the cell cycle and promote apoptosis by inhibiting PI3K/mTOR. The present investigation is the first mechanistic study of NEMF, and these findings provide a new direction for molecular therapy of OV.

In summary, the present results revealed that NEMF is downregulated in OV and can inhibit the proliferation and migration of OV cells. NEMF also inhibits the cell cycle and promotes apoptosis of OV cells by inhibiting PI3K/mTOR, thereby impairing the malignant biological behavior of OV cells. This evidence may provide new insight into the prognostic evaluation and molecular therapy of OV.

## Materials and Methods

### Subcellular localization analysis

We utilized the HPA(Human Protein Atlas,http://www.proteinatlas.org) database as a resource to investigate the subcellular localization of key genes in OV cells(A et al, 2018). Differentially expressed genes (DEGs) and protein–protein interactions (PPIs) were also identified.

In this study, we screened DEGs between tumor tissue samples from TCGA(The Cancer Genome Atlas, https://www.cancer.gov/ccg/research/genome-sequencing/tcga) database using the DESeq2, edgeR, and limma software packages. In addition, we created heatmap and volcano map visualizations using the Pheatmap R package(Ritchie et al, 2015). The PPIs for all common DEGs were analyzed using the search tool for retrieval of interacting genes (STRING) database(Shannon et al, 2003).

### Functional evaluation

We queried the biological processes annotated in the Gene Ontology (GO) and the Kyoto Encyclopedia of Genes and Genomes (KEGG) databases. The adjusted *P*-value threshold for rejecting the null hypothesis was set at 0.05. Explains the most important eight GO terms and the KEGG pathway.

### Study on the infiltration pattern of immune cells

Using the TIMER(Tumor Immune Estimation Resource,http://timer.comp-genomics.or g/) database, we delved into the distribution patterns and functional contributions of various immune subpopulations in the tumor microenvironment, including B cells, two T-cell subtypes (CD4+ and CD8+), neutrophils, macrophages, and dendritic cells.

### Cell culture and reagents

Two human OV cell lines were used, namely OVCAR3 and SKOV3. The human normal ovarian epithelial cell line IOSE80 was also used. OV cell lines were cultured in RPMI 1640 Medium (Vivacell, Shanghai, China) containing 10% fetal bovine serum (Nobimpex, Herbolzheim, Germany) at 37°C in a 5% CO_2_ atmosphere. Culture dishes were purchased from Jet Biofil (Guangzhou, China).

### Reverse transcription and quantitative real-time polymerase chain reaction (RT-qPCR)

Total RNA from OVCAR3, SKOV3, and IOSE80 cells was extracted using the Total RNA Isolation Kit (Foregene, Chengdu, China). Using the Evo M-MLV RT Mix Kit with gDNA Clean for quantitative PCR (AG, Hangzhou, China), mRNA was converted into cDNA, which was used to amplify target genes through quantitative PCR. Quantitative real-time PCR was performed using the CFX96 Real-Time System (Bio-Rad). The following primer sequences of glyceraldehyde-3-phosphate dehydrogenase (GAPDH) were used: GAPDH-forward: 5′-GGACCTGACCTGCCGTCTAG-3′, GAPDH-reverse: 5′-GTAGCCCAGGAT GCCCTTGA-3′. The following primer sequences of NEMF were used: NEMF-forward: 5′-TACTTG ACACCAGGAGAACCC-3′, NEMF-reverse: 5′-CCAAGCAGCACTGTAGCAA A-3′.

### Western blotting

The cells were lysed and the protein concentration was determined using the BCA protein assay kit (Tiangen Biotechnology Co., Ltd., Beijing, China). Proteins were separated using sodium dodecyl sulfate-polyacrylamide gel electrophoresis and transferred onto polyvinylidene difluoride membranes. Finally, a high-sensitivity chemiluminescent reagent (Vazyme, Nanjing, China) was used to visualize specific proteins. The following antibodies were purchased from Proteintech (Wuhan, China): NEMF (11840- 1-AP; 1:1,000), E-cadherin (20874-1-AP; 1:5,000), N-cadherin (22018-1-AP; 1:5,000), vimentin (10366-1-AP; 1:5,000), β-actin (60008-1-Ig; 1:5,000), GAPDH (60004- 1-Ig; 1:5,000), cyclin D1 (CCND1; 26939-1-AP; 1:1,000), P21 (10355-1-AP; 1:1,000), B-cell lymphoma 2 (BCL-2; 60178-1-Ig; 1:1,000), BCL-2-associated X (BAX; 50599-2-Ig; 1:1,000), and mTOR (66888-1-Ig; 1:1,000). Antibodies against phosphorylated-PI3K (p-PI3K; 17366S; 1:1,000), PI3K (4292S; 1:1,000), and p-mTOR (5536S; 1:1,000) were purchased from Cell Signaling Technology (Danvers, MA, USA). The antibody against tubulin (AP0064; 1:5,000) was obtained from Bioworld (Nanjing, China).

### Tissue samples and immunohistochemical staining

Fifty human OV tissue specimens and five normal ovarian tissues were obtained from the Third Affiliated Hospital of Southern Medical University (Guangzhou, China). The use of clinical specimens was approved by the ethics committee of the Third Affiliated Hospital of Southern Medical University. Streptavidin/Peroxidase and 3,3’-diaminobenzidine (DAB) kits (purchased from ZSGB BIO, Beijing, China) were used for immunohistochemical staining.

Immunohistochemical staining was performed according to the instructions provided by the manufacturer. The score was determined by multiplying the score of staining intensity by the score of the proportion of positive cells (0–12 points). Proportion score: Positive cells <25% (1 point), 25–50% (2 points), 50–75% (3 points), >75% (4 points). Strength score: − (0 points), + (1 points), ++ (2 points), +++ (3 points). Total scores of 0–6 and 7–12 indicate low and high expression, respectively. Staining scores were calculated with the assistance of two senior doctors in the Department of Pathology.

### Lentivirus infection

The full-length sequences of NEMF and FLAG-Tag were inserted into the lentivirus (Genechem, Shanghai, China), which contains green fluorescent protein (GFP). To construct cell lines with stable overexpression (OE) of NEMF, OVCAR3 and SKOV3 cells were infected with GFP empty vector or full-length NEMF GFP vector.

### Cell Counting Kit-8 (CCK-8) assay

Cells (2,000 cells/well) were seeded in 96-well plates, and quadruplicate wells were prepared for each condition. CCK-8 reagent (Vazyme) was added every 24 h for 5 days. Two hours after treatment with CCK8 reagent, optical density in each well was measured at 450 nm wavelength.

### 5-Ethynyl-2’-deoxyuridine (EdU) staining

Cells (6,000 cells/well) were seeded in 96-well plates, and triplicate wells were prepared for each condition. Cells were incubated with EdU reagent (10 μM) at 37°C for 2 h and subsequently fixed in 4% paraformaldehyde. EdU staining was performed according to the instructions provided by the manufacturer (Cell-Light EdU Apollo567 *in vitro* kit; Ribobio, Guangzhou, China).

### Colony formation assay

Cells (500 cells/well) were seeded in six-well plates. The cells were incubated at 37°C in a 5% CO_2_ atmosphere for 1 week. Thereafter, the colonies were gently washed twice with phosphate-buffered saline (PBS), fixed in paraformaldehyde, and stained using crystal violet.

### Wound-healing assay

The transfected OV cells were seeded in six-well plates when they reached 95% confluency. When cells were full, a scratch in each well was performed using a 100 μl pipette tip, and the wells were washed thrice with PBS to remove exfoliated cells. Cells were cultured using serum-free medium. Images were captured with a microscope at regular intervals.

### Ghrocyclic peptide staining experiments

1×10^5^ cells were placed into the dish. Following adherence, the cells were fixed with 4% paraformaldehyde, and 0.5% Triton X-100 solution was used to permeate the cells. After washing with PBS, ghrocyclide (200 μl) was added to each dish. Following incubation at room temperature for 1 h, DAPI was added to leave overnight.

### Cell cycle analysis

After transfection, a total of 106 cells were digested and centrifuged. Subsequently, the cells were resuspended with 70% ethanol and left overnight at 4°C. Following treatment of the cells at 37°C and 4°C for 30 min each with the Cell Cycle Assay Kit (DojinDo, Shanghai, China), the DNA content was measured using a FACS bore flow cytometer (BD Biosciences).

### Data Availability

Further information on datasets, materials and protocols should be directed to

## Acknowledgements

This study was supported by the President Foundation of The Third Affiliated Hospital of Southern Medical University(YL202207),the Open Fund of Guangdong Provincial Key Laboratory of Research and Development in Traditional Chinese Medicine (No.KFKT02-005), the Special Research Platform Project of Traditional Chinese Medicine Bureau of Guangdong Province(No.20233005), and the 74th batch of grant from China Postdoctoral Science Foundation(Certificate Number:2023M741570).

## Author Contributions

Q Wang :Data Curation;Formal Analysis;Methodology;Validation;Writing–Original Draft Preparation.

Y Zhu :Data Curation;Investigation;Software;Writing–Original Draft Preparation. Y Zhang :Formal Analysis; Project Administration;Resources;Supervision.

C Chen :Formal Analysis;Investigation;Project Administration;Writing–Original Draft Preparation. A Yin :Investigation;Resources;Supervision.

J Fan :Investigation;Software;Validation.

S Guo :Conceptualization;Funding Acquisition;Methodology; Writing–Review & Editing. Y Liu:Funding Acquisition;Supervision;Writing–Review & Editing.

## Abbreviations

OV: Ovarian cancer
DEG: Differentially expressed genes
GFP: Green fluorescent protein
GO: Gene Ontology
KEGG: Kyoto Encyclopedia of Genes and Genomes
BP: Biological process
CC: Cell composition
MF: Molecular function
NC: Negative control
NEMF: Nuclear export mediator factor
PBS: Phosphate-buffered saline
PCR: Polymerase chain reaction
RQC: Ribosome quality control complex
STRING: Search tool for retrieval of interacting genes
TCGA: The Cancer Genome Atlas
TIMER: Tumor Immune Estimation Resource
FIGO: International Federation of Gynecology and Obstetrics
CI: Confidence interval
HR: Hazard ratio
LTN1: Listerine E3 ubiquitin protein ligase 1
PPI: Protein–protein interaction
TCF25: Transcription factor 25
MDSC: Myeloid-derived suppressor cells

## References

1. A A, M W, G B, R M, R Z, M A, M N, Mt A, W E, A A et al (2021) Biallelic loss-of-function variants in NEMF cause central nervous system impairment and axonal polyneuropathy. Hum Genet 140. doi:10.1007/s00439-020-02226-3.

2. A A, T S, Mc D, T K, C N, Dg E, Ej C, Naj R (2022) The prevalence of mismatch repair deficiency in ovarian cancer: a systematic review and meta-analysis. Int J Cancer 151. doi:10.1002/ijc.34165.

3. A M, Dc L, Y A, C P, Hp K (2018) Maftools: efficient and comprehensive analysis of somatic variants in cancer. Genome research 28. doi:10.1101/gr.239244.118.

4. Akter S, Rahman MA, Hasan MN, Akhter H, Noor P, Islam R, Shin Y, Rahman MDH, Gazi MS, Huda MN et al (2022) Recent advances in ovarian cancer: therapeutic strategies, potential biomarkers, and technological improvements. Cells 11: 650. doi:10.3390/cells11040650.

5. Bengtson MH, Joazeiro CAP (2010) Role of a ribosome-associated E3 ubiquitin ligase in protein quality control. Nature 467: 470–473. doi:10.1038/nature09371.

6. Cao F, Xia W, Dai S, Wang C, Shi R, Yang Y, Guo C, Xu XL, Luo J (2023) Berberine: an inspiring resource for the treatment of colorectal diseases. Biomed Pharmacother = Biomed Pharmacother 167: 115571. doi:10.1016/j.biopha.2023.115571.

7. Chen J, Wei Z, Fu K, Duan Y, Zhang M, Li K, Guo T, Yin R (2022) Non-apoptotic cell death in ovarian cancer: treatment, resistance and prognosis. Biomed Pharmacother = Biomed Pharmacother 150: 112929. doi:10.1016/j.biopha.2022.112929.

8. Chen S, Tang Y, Li Y, Huang M, Ma X, Wang L, Wu Y, Wang Y, Fan W, Hou S (2023) Design and application of prodrug fluorescent probes for the detection of ovarian cancer cells and release of anticancer drug. Biosens Bioelectron 236: 115401. doi:10.1016/j.bios.2023.115401.

9. Crowe-McAuliffe C, Takada H, Murina V, Polte C, Kasvandik S, Tenson T, Ignatova Z, Atkinson GC, Wilson DN, Hauryliuk V (2021) Structural basis for bacterial ribosome-associated quality control by RqcH and RqcP. Mol Cell 81: 115–126.e7. doi:10.1016/j.molcel.2020.11.002.

10. Del Bufalo D, Damia G (2024) Overview of BH3 mimetics in ovarian cancer. Cancer Treat Rev 129: 102771. doi:10.1016/j.ctrv.2024.102771.

11. Dh G, Wx Z (2013) [recent advances on ovarian epithelial cancer: definition, subtypes and pathologic features]. Zhonghua bing li xue za zhi = Chin j pathol 42.

12. Duska LR, Kohn EC (2017) The new classifications of ovarian, fallopian tube, and primary peritoneal cancer and their clinical implications. Ann Oncol: Off J Eur Soc Med Oncol 28: viii8– viii12. doi:10.1093/annonc/mdx445.

13. Ediriweera MK, Tennekoon KH, Samarakoon SR (2019) Role of the PI3K/AKT/mTOR signaling pathway in ovarian cancer: biological and therapeutic significance. Semin Cancer Biol 59: 147–160. doi:10.1016/j.semcancer.2019.05.012.

14. Ersahin T, Tuncbag N, Cetin-Atalay R (2015) The PI3K/AKT/mTOR interactive pathway. Mol Biosyst 11: 1946–1954. doi:10.1039/c5mb00101c.

15. F B, M L, H S, J F, Rl S, I S, A J (2024) Global cancer statistics 2022: GLOBOCAN estimates of incidence and mortality worldwide for 36 cancers in 185 countries. CA: a cancer journal for clinicians 74. doi:10.3322/caac.21834.

16. Gasparri ML, Bardhi E, Ruscito I, Papadia A, Farooqi AA, Marchetti C, Bogani G, Ceccacci I, Mueller MD, Benedetti Panici P (2017) PI3K/AKT/mTOR Pathway in Ovarian Cancer Treatment: Are We on the Right Track? Geburtshilfe Frauenheilkd 77: 1095–1103. doi:10.1055/s-0043-118907.

17. Glaviano A, Foo ASC, Lam HY, Yap KCH, Jacot W, Jones RH, Eng H, Nair MG, Makvandi P, Geoerger B et al (2023) PI3K/AKT/mTOR signaling transduction pathway and targeted therapies in cancer. Mol Cancer 22: 138. doi:10.1186/s12943-023-01827-6.

18. Godbole N, Quinn A, Carrion F, Pelosi E, Salomon C (2023) Extracellular vesicles as a potential delivery platform for CRISPR-cas based therapy in epithelial ovarian cancer. Semin Cancer Biol 96: 64–81. doi:10.1016/j.semcancer.2023.10.002.

19. Hickey KL, Dickson K, Cogan JZ, Replogle JM, Schoof M, D’Orazio KN, Sinha NK, Hussmann JA, Jost M, Frost A et al (2020) GIGYF2 and 4EHP inhibit translation initiation of defective messenger RNAs to assist ribosome-associated quality control. Mol Cell 79: 950–962.e6. doi:10.1016/j.molcel.2020.07.007.

20. Huang J, Chen L, Wu J, Ai D, Zhang J-Q, Chen T-G, Wang L (2022) Targeting the PI3K/AKT/mTOR Signaling Pathway in the Treatment of Human Diseases: Current Status, Trends, and Solutions. J Med Chem 65: 16033–16061. doi:10.1021/acs.jmedchem.2c01070.

21. Huang T-T, Lampert EJ, Coots C, Lee J-M (2020) Targeting the PI3K pathway and DNA damage response as a therapeutic strategy in ovarian cancer. Cancer Treat Rev 86: 102021. doi:10.1016/j.ctrv.2020.102021.

22. Jd C, Al F, C H, Gp D, Sl B, Cm R, G H, S B, Ty M, S S et al (2007) A transforming mutation in the pleckstrin homology domain of AKT1 in cancer. Nature 448. doi:10.1038/nature05933.

23. Jiang Y, Song L, Lin Y, Nowialis P, Gao Q, Li T, Li B, Mao X, Song Q, Xing C et al (2023) ROS-mediated SRMS activation confers platinum resistance in ovarian cancer. Oncogene 42: 1672–1684. doi:10.1038/s41388-023-02679-6.

24. K E (2022) Cell cycle regulation: p53-p21-RB signaling. Cell Death Differ 29. doi:10.1038/s41418-022-00988-z.

25. Kaloni D, Diepstraten ST, Strasser A, Kelly GL (2023) BCL-2 protein family: attractive targets for cancer therapy. Apoptosis: Int J Program Cell Death 28: 20–38. doi:10.1007/s10495-022-01780-7.

26. Knudsen ES, Kumarasamy V, Nambiar R, Pearson JD, Vail P, Rosenheck H, Wang J, Eng K, Bremner R, Schramek D et al (2022) CDK/cyclin dependencies define extreme cancer cell-cycle heterogeneity and collateral vulnerabilities. Cell Rep 38: 110448. doi:10.1016/j.celrep.2022.110448.

27. L K, Sr G (2020) Treatment of epithelial ovarian cancer. BMJ (Clin res ed,) 371. doi:10.1136/bmj.m3773.

28. L Y, J W, P L (2022) Attacking the PI3K/akt/mTOR signaling pathway for targeted therapeutic treatment in human cancer. Semin Cancer Biol 85. doi:10.1016/j.semcancer.2021.06.019.

29. Lheureux S, Braunstein M, Oza AM (2019) Epithelial ovarian cancer: evolution of management in the era of precision medicine. CA Cancer J Clin 69: 280–304. doi:10.3322/caac.21559.

30. Liu C, Zheng W (2011) [advances in origin and pathogenesis of epithelial ovarian cancer]. Zhonghua Bing Li Xue Za Zhi = Chin J Pathol 40: 569–572.

31. M D, C LV, P B, P B, F L, E N, M M (2022) European cancer mortality predictions for the year 2022 with focus on ovarian cancer. Ann oncol : off j Eur Soc Med Oncol 33. doi:10.1016/j.annonc.2021.12.007.

32. Miras I, Estévez-García P, Muñoz-Galván S (2024) Clinical and molecular features of platinum resistance in ovarian cancer. Crit Rev Oncol Hematol 201: 104434. doi:10.1016/j.critrevonc.2024.104434.

33. N G, B A, Ow M, V KM, Eh C, E G (2023) Chemical modulation of cytosolic BAX homodimer potentiates BAX activation and apoptosis. Nat Commun 14. doi:10.1038/s41467-023-44084-3.

34. Nikolatou K, Sandilands E, Román-Fernández A, Cumming EM, Freckmann E, Lilla S, Buetow L, McGarry L, Neilson M, Shaw R et al (2023) PTEN deficiency exposes a requirement for an ARF GTPase module for integrin-dependent invasion in ovarian cancer. EMBO J 42: e113987. doi:10.15252/embj.2023113987.

35. Nunes M, Bartosch C, Abreu MH, Richardson A, Almeida R, Ricardo S (2024) Deciphering the molecular mechanisms behind drug resistance in ovarian cancer to unlock efficient treatment options. Cells 13: 786. doi:10.3390/cells13090786.

36. Occhiuzzi MA, Lico G, Ioele G, De Luca M, Garofalo A, Grande F (2023) Recent advances in PI3K/PKB/mTOR inhibitors as new anticancer agents. Eur J Med Chem 246: 114971. doi:10.1016/j.ejmech.2022.114971.

37. Pa K, Ua M (2023) Clinical and translational advances in ovarian cancer therapy. Nat Cancer 4. doi:10.1038/s43018-023-00617-9.

38. Pb M, Y K-T, Rb S, G R, Je S, R K, R Y, T M, C G, W A et al (2020) NEMF mutations that impair ribosome-associated quality control are associated with neuromuscular disease. Nature communications 11. doi:10.1038/s41467-020-18327-6.

39. Ritchie ME, Phipson B, Wu D, Hu Y, Law CW, Shi W, Smyth GK (2015) limma powers differential expression analyses for RNA-sequencing and microarray studies. Nucleic Acids Res 43: e47. doi:10.1093/nar/gkv007.

40. Shannon P, Markiel A, Ozier O, Baliga NS, Wang JT, Ramage D, Amin N, Schwikowski B, Ideker T (2003) Cytoscape: a software environment for integrated models of biomolecular interaction networks. Genome Res 13: 2498–2504. doi:10.1101/gr.1239303.

41. Shao S, Brown A, Santhanam B, Hegde RS (2015) Structure and assembly pathway of the ribosome quality control complex. Mol Cell 57: 433–444. doi:10.1016/j.molcel.2014.12.015.

42. Siegel RL, Miller KD, Fuchs HE, Jemal A (2021) Cancer statistics, 2021. CA Cancer J Clin 71: 7–33. doi:10.3322/caac.21654.

43. Solidoro R, Centonze A, Miciaccia M, Baldelli OM, Armenise D, Ferorelli S, Perrone MG, Scilimati A (2024) Fluorescent imaging probes for in vivo ovarian cancer targeted detection and surgery. Med Res Rev 44: 1800–1866. doi:10.1002/med.22027.

44. Stamp JP, Gilks CB, Wesseling M, Eshragh S, Ceballos K, Anglesio MS, Kwon JS, Tone A, Huntsman DG, Carey MS (2016) BAF250a expression in atypical endometriosis and endometriosis-associated ovarian cancer. Int J Gynecol Cancer: Off J Int Gynecol Cancer Soc 26: 825–832. doi:10.1097/IGC.0000000000000698.

45. Sun Y, Liu Y, Ma X, Hu H (2021) The influence of cell cycle regulation on chemotherapy. Int J Mol Sci 22: 6923. doi:10.3390/ijms22136923.

46. T U, M S, T O, S A, T N, T N, A M, T I (2021) Failure to degrade CAT-tailed proteins disrupts neuronal morphogenesis and cell survival. Cell Rep 34. doi:10.1016/j.celrep.2020.108599.

47. Tewari D, Patni P, Bishayee A, Sah AN, Bishayee A (2022) Natural products targeting the PI3K-akt-mTOR signaling pathway in cancer: a novel therapeutic strategy. Semin Cancer Biol 80: 1–17. doi:10.1016/j.semcancer.2019.12.008.

48. Thrun A, Garzia A, Kigoshi-Tansho Y, Patil PR, Umbaugh CS, Dallinger T, Liu J, Kreger S, Patrizi A, Cox GA et al (2021) Convergence of mammalian RQC and C-end rule proteolytic pathways via alanine tailing. Mol Cell 81: 2112–2122.e7. doi:10.1016/j.molcel.2021.03.004.

49. Vaughan S, Coward JI, Bast RC, Berchuck A, Berek JS, Brenton JD, Coukos G, Crum CC, Drapkin R, Etemadmoghadam D et al (2011) Rethinking ovarian cancer: recommendations for improving outcomes. Nat Rev Cancer 11: 719–725. doi:10.1038/nrc3144.

50. X B, T J, F A, H L, B S, Da H, Ma M (2005) Drosophila caliban, a nuclear export mediator, can function as a tumor suppressor in human lung cancer cells. Oncogene 24. doi:10.1038/sj.onc.1208962.

51. X Y, M P, J Y, Ca H, Ss T, J L, B L (2024) Quantifiable TCR repertoire changes in prediagnostic blood specimens among patients with high-grade ovarian cancer. *Cell rep*, Med 5. doi:10.1016/j.xcrm.2024.101612.

52. Y X, M B, H G, M L (2022) Multi-omics approaches for biomarker discovery in early ovarian cancer diagnosis. EBioMedicine 79. doi:10.1016/j.ebiom.2022.104001.

